# The methodological quality of animal studies: A cross-sectional study based on the SYRCLE’s risk of bias tool

**DOI:** 10.1101/701110

**Authors:** Weiyi Zhang, Yanbiao Jiang, Zhizhong Shang, Nan Zhang, Gongcai Tao, Ting Zhang, Kaiyan Hu, Yanfei Li, Xiue Shi, Yanying Zhang, Jiao Yang, Bin Ma, Kehu Yang

**Affiliations:** Evidence Based Social Science Research Center, School of Public Health, Lanzhou University, Lanzhou, China; Key Laboratory of Evidence Based Medicine and Knowledge Translation of Gansu Province, Lanzhou, China; The Second Clinical Medical School, Lanzhou University, Lanzhou, China; The First Clinical Medical School, Lanzhou University, Lanzhou, China; Evidence Based Medicine Center, School of Basic Medical Sciences, Lanzhou University, Lanzhou, China; Evidence Based Nursing Center, School of Nursing, Lanzhou University, Lanzhou, China; Institute for Evidence Based Rehabilitation Medicine of Gansu Province, Lanzhou, China; Gansu University of Traditional Chinese Medicine, Lanzhou, China

**Keywords:** Animal experiments, SYRCLE’s risk of bias tool, Methodological quality

## Abstract

**Objective:** To assess the methodological quality of animal studies published in China and abroad using the SYRCLE’s risk of bias tool, and to provide references to improve the methodological quality of animal studies to encourage high quality preclinical studies.

**Methods:** An electronic search was performed in the Chinese Scientific Citation Database (CSCD) and Web of Science from 2014 to October 2018. Document screening and data extraction were performed independently by four researchers. The methodological quality of the included studies was assessed using the SYRCLE’s risk of bias tool. Statistical analysis was performed using SPSS23.

**Results:** A total of 2764 animal studies were included. Of the studies, 984 were published in English and 1780 were in Chinese. The citation frequency of more than 90% of the included studies was less than 5. The results of methodological quality assessment showed that 36.36% (8/22) of the sub-items were rated as “low risk” in more than 50% of the included studies, of which 75% (6/8) were rated as “low risk” in more than 80% of the included studies. A total of 59.09% (13/22) of the sub-items were rated as “low risk” in less than 30% of the included studies, of which 92.31% (12/13) were rated as “low risk” in less than 10% of the included studies. The incidence of “low risk” Chinese studies regarding performance bias, detection bias and reporting bias were lower than English studies. For foreign studies, more attention should be paid to selection bias, attrition bias, and reporting bias.

**Conclusion:** We identified limitations in the methodological quality of animal experiment studies published in China and abroad. We therefore suggest that it is necessary to take targeted measures to popularize the SYRCLE’s risk of bias tool to effectively improve the design and implementation of animal experiments, and guide study development.

## Introduction

Animal experiments are key to preclinical research. As a bridge between basic science and clinical research, the quality of animal experiments dictates the success of many research fields.[1, 2] And experiment quality is dependent upon scientific design and standard implementation. According to the National Centre for the 3Rs (The National Centre for the Replacement, Refinement and Reduction of Animals in Research), many animal experiments funded by 3Rs had problems in method design. Up to 87% and 86% of experiments fail to perform “random allocation” or “blind method”.[3] In addition, in animal experiments where the principle of “randomization” was implemented, only 9% of studies described specific methods of randomization, which led to low conversion and utilization rates. To improve the methodological quality of animal experiments and promote the transparency of the research process, a number of scholars of the SYRCLE Center (SYstematic Review Center for Laboratory animal Experimentation) including Hooijmans and colleagues, drafted the SYRCLE’s risk assessment tool (SYRCLE’s risk of bias tool for animal studies) in 2014.[2] This tool highlighted that the design and effective implementation of animal experiments could be improved.[4] Since its publication, some domestic scholars have analyzed it[5] and there have been study researching the methodological quality of animal experiments based on this tool, but it was limited to the incidence of stroke, and lack representation from other fields.[6] In addition, a horizontal comparison of the methodological quality of animal experiments published at home and abroad has not been performed.

In this study, we performed a comprehensive analysis of animal experiments published in China and abroad to assess the inherent risk of bias based on SYRCLE’s risk of bias tool. We analyzed and discussed the differences between Chinese and English studies regarding method design, and highlighted possible causes of bias to provide references for the design and implementation of future animal experiments.

## 1. Material and methods

### 1.1 Inclusion and exclusion criteria

All interventional animal experiments were included without restrictions on the type of intervention or animal species. The exclusion criteria were as follows: (1) Self-controlled studies; (2) non-medical animal experiments; (3) In vitro experiments based on animal tissues, organs or cells; (4) experiments including both animals and humans; (5) studies on molecular mechanisms; (6) studies published in duplicate.

### 1.2 Search strategy

To compare the quantity and quality of domestic and foreign studies, animal experiments were identified through searching the Chinese Science Citation Database (CSCD) and Web of Science. Documents included in the CSCD database were from highest-ranking journals in China, and so were equivalent to studies collected by the Science Citation Index (SCI). As the SYRCLE’s risk of bias tool was published in 2014, the search time was limited from 2014 to October 2018. The following search terms were used: animal experimentation, animal experiments, animal experiment, animal study, animal studies, animal research, in vivo, in vivo study, mice, mus, mouse, murine, rats, rat, pigs, pig, swine, swines, guinea pigs, guinea pig, cavia, rabbits, rabbit, dogs, dog, canine, canines, canis, sheep, goats, goat, monkey, monkeys, ape, apes, orangutan, paniscus, pan paniscus, bonobo, bonobos, pan troglodytes, chimpanzee, chimpanzees, gorilla, gorillas, pongo, frog, frogs, toad, toads.

### 1.3 Screening and data extraction

Four researchers (Gongcai Tao, Nan Zhang, Yanbiao Jiang, Zhizhong Shang) were trained to master the inclusion and exclusion criteria of this study and the items of the SYRCLE’s risk of bias tool. Approximately 10% of the samples were randomly selected for pre-experiments and Kappa values were calculated (Kappa value= 0.72). Finally, the researchers screened the documents and extracted the data independently according to pre-designed data extraction table and cross-checked in pairs. Disagreements were decided by a third party (Bin Ma). Data extraction included: (1) general characteristics of the included studies: author, year of publication, country, times cited, animal species, sample size and interventions; (2) risk of bias assessments: coincidence rates of “low risk” for 22 sub-items.

### 1.4 Statistical analysis

Statistical analysis was performed using SPSS23. Counting data were statistically described by numbers and percentages (%), and measurement data were described by the median and interquartile range. Chi-square tests were used for group comparisons. A p-value less than 0.05 was considered statistically significant.

## 2. Results

### 2.1 Screening procedures

A total of 33264 related documents were obtained from the initial search, of which 18260 were published in Chinese and 15004 were published in English. After the removal of duplicate studies and those failing to meet the inclusion criteria, 2764 animal experiments were finally included (1780 in Chinese and 984 in English). (See Fig 1)

**Figure 1.**
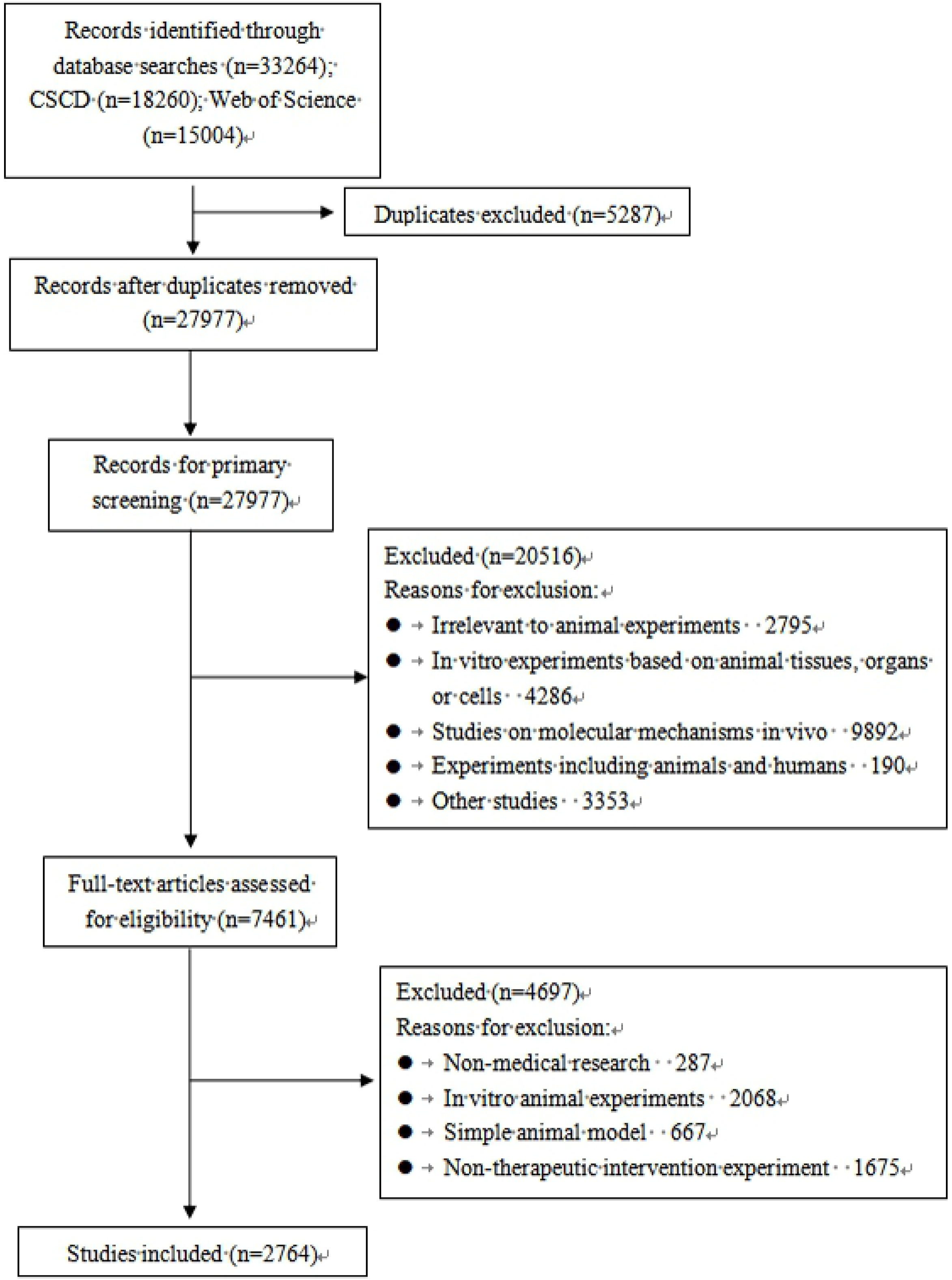
Flow diagram of study selection process.

### 2.2 Study characteristics

The median sample size of the Chinese studies was 50, and the interquartile range was 32 to 70. For English studies, the median sample size was 36 and the interquartile range was 24 to 51. The citation frequency of more than half (52.89%, 1462/2764) of the studies was zero. The percentage of Chinese studies (67.30%, 1198/1780) with zero citations was significantly higher than those of English studies (26.83%, 264/984, p<0.001). The most commonly used animal species were rats (87.99%, 2432/2764), rabbits (6.22%, 172/2764), and pigs (1.48%, 41/2764). Amongst them, the differences between Chinese and English studies were statistically significant for rats (p<0.001) and sheep (p=0.002). The top three interventions were: drug interventions (74.17%, 2050/2764), surgical interventions (7.56%, 209/2764), and behavioral-dietary interventions (4.59%, 127/2764). Amongst them, the differences between Chinese and English studies were statistically significant regarding the choice of drug (p<0.001) and acupuncture intervention (p<0.001). (See Figs 2-4)

**Figure 2.**
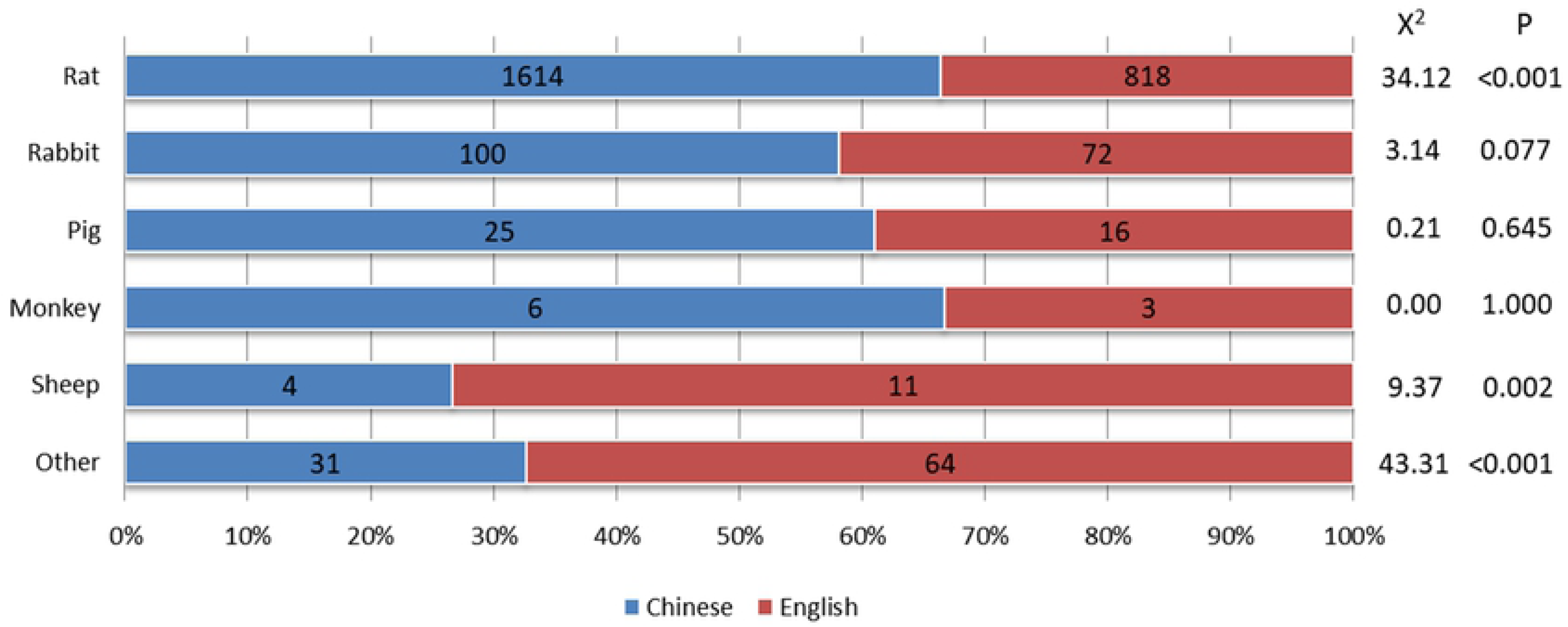
Animal species of included studies.

**Figure 3.**
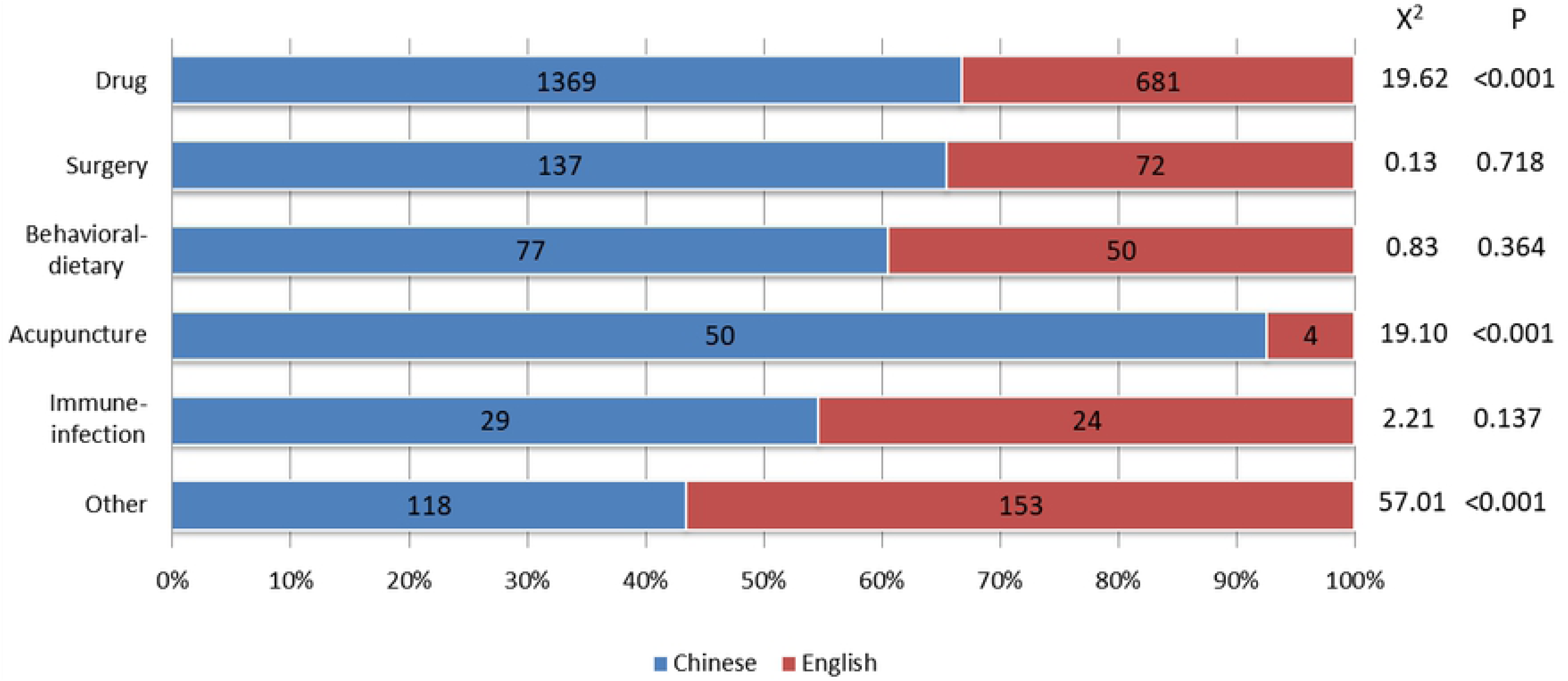
Interventions of included studies.

**Figure 4.**
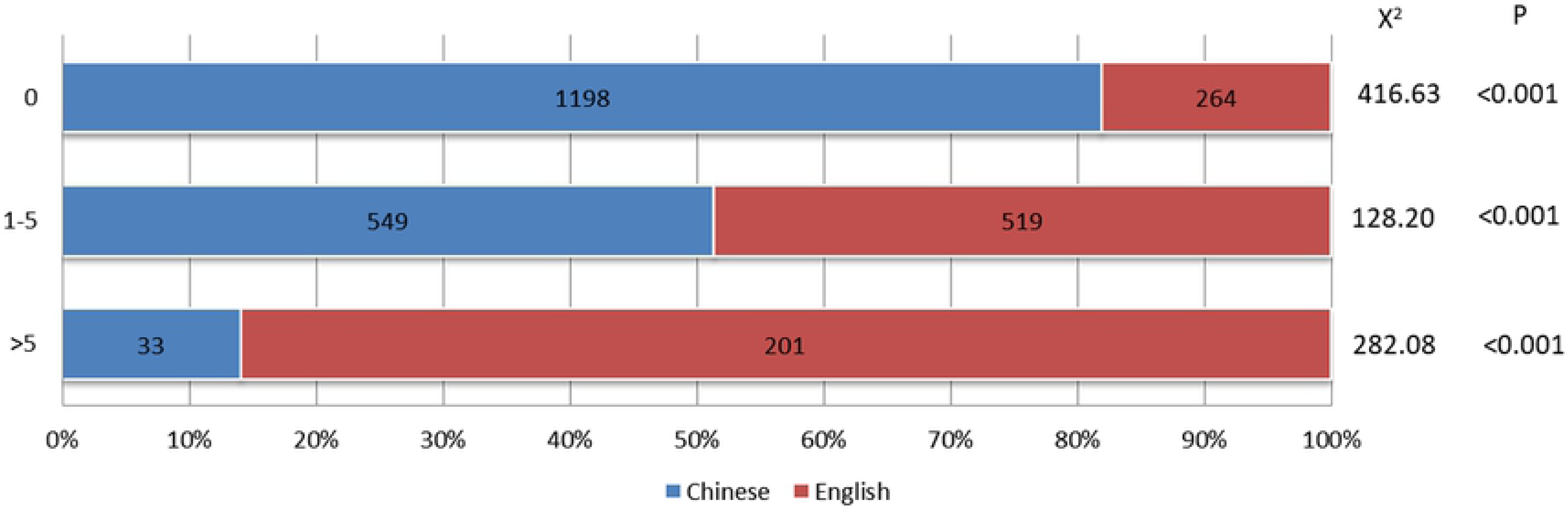
Citation frequency of included studies.

### 2.3 Assessment of the risk of bias

The quality of the included animal experiments was assessed using the SYRCLE’s risk of bias tool that contains 10 entries and 22 sub-items. A total of 36.36% (8/22) of the sub-items were rated as “low risk” in more than 50% of the included studies, of which 75% (6/8) were rated as “low risk” in more than 80% of the included studies. A total of 59.09% (13/22) of the sub-items were rated as “low risk” in less than 30% of the included studies, of which 92.31% (12/13) were rated as “low risk” in less than 10% of the included studies. (See Table 1)

**Table 1.** Risk of bias in the included studies.

| Type of bias | Domain | Item | Sub-item | "Low risk" item (total) |  | "Low risk" item (Chinese) |  | "Low risk" item (English) |  | Chi-square value | P value |
| --- | --- | --- | --- | --- | --- | --- | --- | --- | --- | --- | --- |
|  |  |  |  | N | % | N | % | N | % |  |  |
| Selection bias | Sequence generation | (1) | 1 | 548 | 19.83 | 395 | 22.19 | 153 | 15.55 | 17.59 | <0.001 |
|  | Baseline characteristics | (2) | 2 | 1769 | 84.80 | 1489 | 83.65 | 855 | 86.89 | 5.16 | 0.02 |
|  |  |  | 3 | 46 | 4.62 | 39 | 13.40 | 7 | 5.43 | 5.83 | 0.02 |
|  |  |  | 4 | 2490 | 92.29 | 1626 | 95.91 | 864 | 91.14 | 2.72 | 0.10 |
|  | Allocation concealment | (3) | 5 | 23 | 0.83 | 15 | 0.84 | 8 | 0.81 | 0.01 | 0.93 |
| Performance bias | Random housing | (4) | 6 | 33 | 1.19 | 3 | 0.17 | 30 | 3.05 | 44.56 | <0.001 |
|  |  |  | 7 | 112 | 4.10 | 3 | 0.17 | 109 | 11.43 | 200.00 | <0.001 |
|  | Caregivers blinding | (5) | 8 | 83 | 3.00 | 39 | 2.19 | 44 | 4.47 | 11.32 | <0.001 |
| Detection bias | Random outcome assessment | (6) | 9 | 961 | 34.77 | 713 | 40.06 | 248 | 25.20 | 61.64 | <0.001 |
|  | Outcome assessor blinding | (7) | 10 | 166 | 6.01 | 92 | 5.17 | 74 | 7.52 | 6.21 | 0.013 |
|  |  |  | 11 | 240 | 9.24 | 26 | 1.54 | 214 | 23.52 | 340.57 | <0.001 |
| Attrition bias | Incomplete outcome data | (8) | 12 | 1743 | 63.06 | 1241 | 69.72 | 502 | 51.02 | 95.16 | <0.001 |
|  |  |  | 13 | 79 | 7.74 | 63 | 11.69 | 16 | 3.32 | 24.96 | <0.001 |
|  |  |  | 14 | 54 | 5.29 | 38 | 7.10 | 16 | 3.32 | 8.05 | 0.005 |
|  |  |  | 15 | 14 | 1.37 | 11 | 2.04 | 3 | 0.62 | 3.79 | 0.052 |
| Reporting bias | Reporting bias | (9) | 16 | 31 | 1.12 | 0 | 0 | 31 | 3.15 | 56.71 | <0.001 |
|  |  |  | 17 | 2715 | 99.20 | 1770 | 99.44 | 945 | 99.16 | 0.73 | 0.392 |
| Other | Other bias | (10) | 18 | 2676 | 96.82 | 1717 | 96.46 | 959 | 97.46 | 2.05 | 0.152 |
|  |  |  | 19 | 2746 | 99.35 | 1770 | 99.44 | 976 | 99.19 | 0.62 | 0.432 |
|  |  |  | 20 | 2752 | 99.57 | 1774 | 99.66 | 978 | 99.39 | 0.55 | 0.458 |
|  |  |  | 21 | 1464 | 52.97 | 926 | 52.02 | 538 | 54.67 | 1.79 | 0.181 |
|  |  |  | 22 | 81 | 7.92 | 23 | 4.27 | 58 | 11.98 | 21.01 | <0.001 |

For the 5 sub-items related to selective bias (sub-items 1 to 5), the coincidence rates of “low risk” for sub-item 3 and sub-item 5 were less than 5% (4.62%; 0.83%). The coincidence rates of “low risk” of sub-item 2 amongst Chinese studies was significantly lower than those of English studies (p= 0.02). The coincidence rate of “low risk” sub-items 1 and 3 amongst Chinese studies were significantly higher than those of English studies (p<0.001; p=0.02).

For the 3 sub-items related to performance bias (sub-items 6 to 8), the coincidence rates of “low risk” were less than 5% (1.19%; 4.10%; and 3.00%, respectively). The coincidence rates of “low risk” of the 3 sub-items amongst Chinese studies were significantly lower than those of English studies (p<0.001).

For the 3 sub-items related to detection bias (sub-items 9 to 11), the coincidence rates of “low risk” for sub-items 10 and 11 were less than 10% (6.01%; 9.24%). The coincidence rates of the “low risk” sub-items amongst Chinese studies were significantly lower than those of English studies (p=0.01; p<0.001). However, the coincidence rates of “low risk” in sub-item 9 amongst Chinese studies was significantly higher than English studies (p<0.001).

For the 4 sub-items related to attrition bias (sub-items 12 to 15), the coincidence rates of “low risk” for sub-items 13, 14 and 15 were all less than 10%. The coincidence rates of “low risk” sub-items 12, 13, and 14 amongst the English studies were significantly lower than Chinese studies (p<0.001; p<0.001; p=0.005, respectively).

For the 2 sub-items related to reporting bias (sub-items 16 to 17), the coincidence rate of “low risk” for sub-item 16 was less than 5% (1.12%), and the coincidence rate of “low risk” amongst Chinese studies was significantly lower than that of English studies (p<0.001).

For the 5 sub-items related to other types of bias (sub-items 18 to 22), only the coincidence rate of “low risk” for sub-item 22 was less than 10% (7.92%), and the coincidence rate of “low risk” amongst Chinese studies was significantly lower than that of English studies (p<0.001).

## 3. Discussion

### 3.1 General characteristics

The results of this study showed that the citations of more than 90% of the included studies were less than 5, and 52.89% of the studies have not been cited. The percentage of Chinese studies with zero citations was significantly higher than English studies (p<0.001). This may be related to the low quality of animal experimental studies published in Chinese, as previously suggested by Shuang et al.[6] The most commonly used animals both at home and abroad were rats, which may be because rats are easy to breed, and inbreeding mode makes individual differences between rats small, which is conducive to the experiments.[7, 8] Regarding interventions, the number of Chinese studies using acupuncture were significantly higher than English studies, which is likely due to its known association with traditional Chinese medicine.[9]

### 3.2 Summary of the methodological quality assessment

The SYRCLE’s risk of bias tool is a globally unique tool that focuses on the assessment of the intrinsic authenticity of animal experiments. It includes six types of bias: (1) selection bias; (2) performance bias; (3) detection bias; (4) attrition bias; (5) reporting bias and (6) other bias.[4]

Selection bias represents systematic errors between the baseline characteristics of comparison groups.[10] Because the sample size of animal experiments is smaller than that of clinical trials, if it is not controlled, an unbalanced baseline in some important characteristics would influence the study results.[2] According to this study, the coincidence rates of “low risk” in all included studies were relatively higher for the “baseline characteristics balance (sub-item 2)” and “time of disease induction (sub-item 4)”, but the coincidence rates of “low risk” when “describing a random component in the sequence generation process (sub-item 1)” was less than 30%. The coincidence rates of “low risk” for the “adjustment of unequal distributions of baseline characteristics (sub-item 3)” and the “allocation concealment (sub-item 5)” were less than 5%. The coincidence rates of domestic studies on sub-item 2 were lower than those of foreign studies (p=0.02). However, the coincidence rates of foreign studies on sub-items 1 and 3 were lower than those of domestic studies (p<0.001). Future studies should therefore pay more attention to the correct implementation of random methods, such as the referral to random number tables or computer random number generators. It is also important to perform allocation concealment through sequentially numbered sealed envelopes.[11] Schulz and colleagues showed that strict randomization without allocation concealment results in an exaggeration of the treatment efficacy by 30% to 41%.[12] Domestic studies should pay more attention to the balance of baseline characteristics, whilst foreign studies should adjust unbalanced distributions of the relevant baseline characteristics.

Performance bias refers to systematic errors in the nursing of animals and exposure factors, in addition to the interventions of interest.[13, 14] Animal experiments differ from clinical trials as the setting can influence the study outcome. As an exemplar, the light intensity in cages stored on the top shelve of animal houses is four times higher than that in the cages at the bottom, which can influence reproduction and animal behavior.[15, 16] The results of this study showed that for the 3 sub-items related to performance bias, the coincidence rates of “low risk” in both Chinese and English studies were low, whilst the coincidence rates of “low risk” Chinese studies were significantly lower than foreign studies (P<0.001). This may be related to the lack of attention to performance bias of the relevant studies in China. Ma et al. showed that only 30.8% to 41.4% of domestic researchers considered it necessary to implement “random housing” and “caregivers and researcher blinding”.[17] Future studies should pay more attention to the influence of performance bias. For example, ID cards of individual animals or cage labels should be coded with the same appearance, and the circumstances should be specified in both experimental and control groups during the intervention.

Detection bias refers to systematic errors in the measurement of outcomes.[2] In animal experiments, although it is not necessary to blind animals, the researchers are also animal breeders, which may lead to subjective bias.[18] In addition, due to the characteristics of animal experiments, most animals have circadian rhythm phenomenon, such as lipid metabolism, altered neurotransmitter levels, and pharmacokinetics which can change the cycle/circadian rhythm.[19-21] If the results are not measured by randomization and are instead measured at a single timepoint, the risk of detection bias increases. Our study showed that for the 3 sub-items related to detection bias, the coincidence rates of “low risk” were less than 50%, of which “outcome assessor blinding (sub-item 10)” and “the outcome is not likely to be influenced by lack of blinding (sub-item 11)” were even less than 10%. The coincidence rates of “low risk” for these two sub-items in Chinese studies were significantly lower than foreign studies (P<0.001), but Chinese studies performed to a higher level in sub-item 9 (P<0.001). Future studies should not only pay attention to random outcome assessment including referral to a random number table or the use of random number generators, but further implementing outcome assessor blinding using the same outcome assessment method in both experimental and control groups.

Attrition bias is due to the loss of data caused by death or loss of animals, which impacts the final datasets.[10] The results of this study showed that for the 4 sub-items related to attrition bias, the coincidence rates of “low risk” of “incomplete outcome data” in studies published in Chinese and English were relatively high (more than 50%). However, future studies should pay attention to the processing of missing data and strictly evaluate the influence of the data on the study outcome. For instance, in clinical trials, the intention-to-treat analysis (ITT analysis) is used to process missing data, which is also applicable for animal experiments.[22-24]

Reporting bias refers to systematic errors between reported and unreported results, which may be related to the easier interpretation and publication of positive results, or the existence of conflicts of interest amongst researchers that influence study outcomes.[25, 26] This study showed that the “low risk” coincidence rates for both domestic and foreign studies on “availability of study protocol and all pre-specified primary and secondary outcomes were reported in the manuscript (sub-item 16)” were low. Although the low-risk coincidence rates of English studies were higher than those of Chinese studies (p<0.001), the rates were as low as 3.15%. For clinical studies, registration with the WHO or other first-level registration platforms are required prior to the inclusion of the first case and relevant journals are encouraged to publish their protocols. No such requirements exist for animal experiments.[27-29] However, the publication of protocols can enhance rigor and reduce bias. For future studies, it is necessary to register standardized protocols and clearly report all expected results.[30] In addition, in terms of other bias, the quality of Chinese studies needs to be improved on “adding new animals to the experiment to replace drop-outs from the original population (sub-item 22)”.

### 3.3 Strengths and limitations

This study was the first to compare and analyze the methodological quality of animal experiments published in Chinese and English based on the SYRCLE’s risk of bias tool. Some limitations do however remain. Firstly, the research samples were obtained from the CSCD database and Web of Science. The results may not represent the methodological quality of animal experiments published in journals that were not included in the two databases. Secondly, the risk of bias assessments were based on literature reports and we did not contact the authors to verify the details of the specific implementation of their experiments.

## 4. Conclusions

In summary, limitations in the methodological quality of animal experimental studies published in Chinese and English remain. For domestic studies, problems in performance bias, detection bias, and reporting bias are prevalent. For foreign studies, more attention should be paid to selection bias, attrition bias, and reporting bias. To effectively improve the design and implementation of animal experiments and guide their development, it is necessary to take targeted measures to promote and popularize the SYRCLE’s risk of bias tool.

## Acknowledgments

We thank the following researchers for the assistance: Peijing Yan and Meixuan Li for their support during the study.

